# SuperSpot: Coarse Graining Spatial Transcriptomic Data into Metaspots

**DOI:** 10.1101/2024.06.21.599998

**Authors:** Matei Teleman, Aurélie AG Gabriel, Léonard Hérault, David Gfeller

## Abstract

**Summary:** Spatial Transcriptomics is revolutionizing our ability to phenotypically characterize complex biological tissues and decipher cellular niches. As of today, thousands of genes can be detected across hundreds of thousands of spots. Akin to standard single-cell RNA-Seq data, spatial transcriptomic data are very sparse due to the limited amount of RNA within each spot. Building upon the metacell concept, we present a workflow, called SuperSpot, to combine adjacent and transcriptionally similar spots into “metaspots”. The process involves representing spots as nodes in a graph with edges connecting spots in spatial proximity and edge weights representing transcriptional similarity. Hierarchical clustering is used to aggregate spots into metaspots at a user-defined resolution. We demonstrate that metaspots can be used to reduce the size of spatial transcriptomic data and remove some of the dropout noise.

**Availability and implementation:** SuperSpot is an R package available at https://github.com/GfellerLab/SuperSpot.

## Introduction

Spatial transcriptomics enables researchers to simultaneously capture the spatial organization of cells within a tissue slice and their gene expression profile. Two main groups of spatial transcriptomic technologies have been developed. The sequencing-based technologies (e.g. 10x Visium) use spatially barcoded probes or oligonucleotide spots to capture mRNA molecules from a tissue section. Following sequencing, the spatial transcriptomic data can be mapped back to the original tissue section (Rodriques *et al*. 2019). The imaging-based technologies (e.g. Nanostring CosMx) rely mainly on fluorescence in situ hybridization or in situ sequencing to detect specific mRNA molecules within a tissue section, followed by high- resolution imaging and computational analysis to map the expression of hundreds to thousands of genes in their native tissue context (Chen *et al*. 2015; He *et al*. 2022; Shi *et al*. 2023). The captured mRNAs are assigned to spots after cell segmentation.

Historically, the sequencing-based approaches were limited to a few thousand spots, with each spot consisting of roughly 10 cells (Du *et al*. 2023; Park *et al*.). More recently, the resolution of these approaches has significantly increased allowing researchers to profile tens of thousands of spots at almost single cell resolution (Nagendran *et al*. 2023). The imaging-based methods can reach sub-cellular resolution and profile hundreds of thousands of spots (Chen *et al*. 2015). Akin to single-cell RNA-Seq (scRNA-Seq) both sequencing-based and imaging-based spatial transcriptomic data are very sparse (from 80% to 95% of zeros) in part because of the limited amount of RNA in each spot (Zhao *et al*. 2022).

To simultaneously address size and sparsity issues in scRNA-seq data, metacells have been proposed, with tools like MetaCell, SuperCell or SEACells developed for this purpose (Baran *et al*. 2019; Ben-Kiki *et al*. 2022; Bilous *et al*. 2022, 2024;; Persad *et al*. 2023). Metacells consist of disjoint groups of cells showing very high transcriptomic similarity which are aggregated together (Bilous *et al*. 2024). Metacell approaches reduce technical noise and computational resource requirements while preserving biological heterogeneity (Baran *et al*. 2019; Ben-Kiki *et al*. 2022; Bilous *et al*. 2022, 2024;; Persad *et al*. 2023). The ratio between the number of cells and the number of metacells is defined as the graining level, typically designated with the Greek letter gamma (γ) (Bilous *et al*. 2024). In many metacell workflows the graining level can be determined by the users which is convenient to explore different values (Ben-Kiki *et al*. 2022; Bilous *et al*. 2022; Persad *et al*. 2023).

In this work, we expand the metacell concept to spatial transcriptomic data with the aim of simultaneously reducing the size and sparsity of such data by merging spots into metaspots and preserving both the spatial and transcriptomic information.

## Results

To adapt the concept of metacells to spatial transcriptomics, we developed the SuperSpot workflow. This workflow builds a graph on the coordinates of the spots and uses transcriptional similarity as weights for the edges (Fig. 1A). For sequencing-based data generated with 10x Visium technology we consider edges between adjacent spots. For imaging-based data we started with a K-Nearest Neighbor (KNN) graph with K=15. To avoid long-range connections between distant spots in sparse regions of the images, the top 40% of connections based on Euclidian distances were removed from the graph. To compute the weights, we performed Principal Component Analysis (PCA) on the normalized count matrix. For each pair of connected spots in the graph, a transcriptomic distance was computed from the top 30 Principal Components space. The resulting distance was converted into a transcriptional similarity measure by the function: 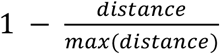. As in the SuperCell method (Bilous *et al*. 2022), the walktrap hierarchical clustering (Pons and Latapy 2005) was then applied on the graph to merge the spots into metaspots. In this way, users can rapidly explore different graining levels (γ) without having to recompute the clustering for each γ. The metaspots are defined transcriptomically by aggregating raw counts and renormalizing them at the metaspot level, and spatially by computing the centroid of the spatial coordinates of the spots they consist of.

**Figure 1:**
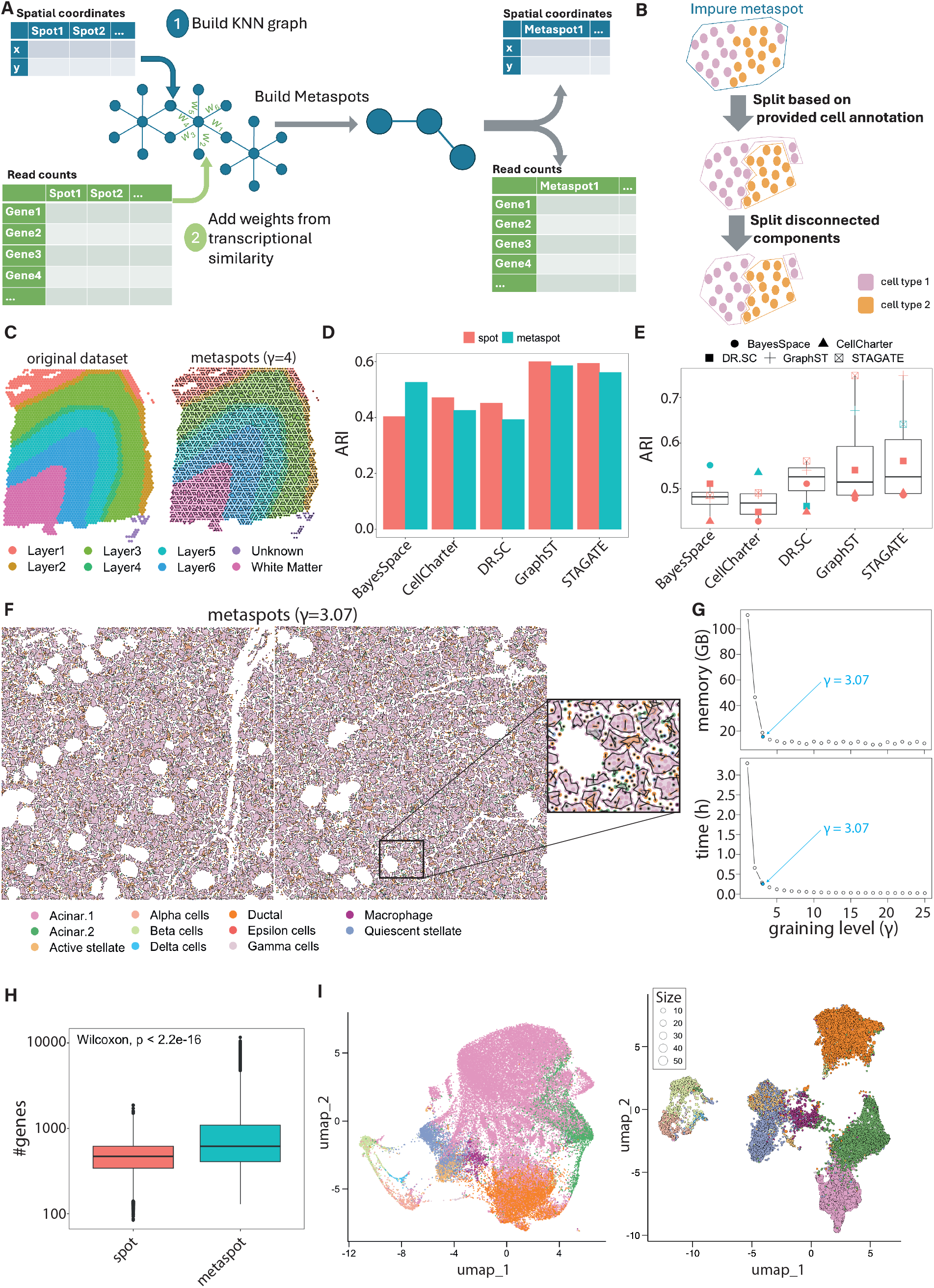
SuperSpot overview and applications. **A)** Illustration of the SuperSpot pipeline to build metaspots in spatial transcriptomic data. **B)** Splitting process of metaspots to guarantee both purity according to some annotation and spatial coherence. **C)** 10x Visium mouse cortex dataset at the spot and metaspot (γ = 4) level. The spots and the metaspots are colored based on the layers of the cortex and white matter. The “Unknown” label corresponds to unannotated spots. **D)** ARI scores of different spatial clustering methods with respect to the brain layer annotation at spot (red) and metaspot (blue) levels. The ARI score is computed as the mean over 10 runs with different seeds for each clustering method. **E)** ARI scores between the clusters computed at the spot and metaspot levels by each clustering method (blue dots) and the clusters computed by different clustering methods at spot level (red dots). The ARI score is computed as the mean over 10 runs with different seeds for each clustering method. **F)** Nanostring CosMx human pancreas dataset containing two slides at the metaspot (γ = 3.07) level. The spots and the metaspots are colored based on the cell types. **G)** Peak memory (GB) and elapse time needed for computing normalization and spatially variable features of Nanostring CosMx human pancreas dataset containing 48,944 spots at the metaspot level as a function of γ. Blue dot represents the amount of time and memory for the split metaspots with the final γ of 3.07. **H)** Boxplot of the number of detected genes per spot (red) and metaspot (blue) for whole CosMx human pancreas dataset. **I)** UMAP visualization of the original CosMx human pancreas dataset at the spot level (left) and at the metaspot level (right). For the UMAP at the metaspot level, the size of the dots is proportional to the number of spots contained in the metaspot. Colors correspond to the legend of panel G.

When a spot categorical annotation (e.g. cell types or niches) is provided, metaspots containing spots with distinct annotations can be split (Fig. 1B). To this end, metaspots are first split into smaller metaspots for each category of the annotation. To keep spatial coherence, we split a second time the resulting metaspots if they consist of disconnected components in the initial KNN graph. These two splitting steps induce a final γ smaller than the initial one.

To evaluate how metaspots can be used for downstream analyses, we investigated whether the results of clustering applied on metaspots are consistent with those obtained at the spot level. We applied different spatial clustering methods on a reference mouse cortex dataset, generated with 10x Visium technology (Maynard *et al*. 2021), including GraphST (Long *et al*. 2023), STAGATE (Dong and Zhang 2022), CellCharter (Varrone *et al*. 2024), DR.SC (Liu *et al*. 2022) and BayesSpace (Zhao *et al*. 2021). This analysis was done both at the spot and metaspot level (with a γ of 4 and without splitting the metaspots based on an annotation) (Fig. 1C). The obtained clusters were compared to brain layer annotation of the spots. By computing the Adjusted Rand Index (ARI) scores between the clusters and the ground truth, we observed similar results at the spot and metaspot resolution (Fig. 1D). Additionally, the differences in the clusters computed at the spot and metaspot levels using the same algorithm were comparable (i.e. similar ARI) to differences obtained when using different clustering algorithms at the spot resolution (Fig. 1E).

We next applied SuperSpot to a publicly available Nanostring CosMx dataset of a human pancreas (CosMx SMI Human Pancreas FFPE Dataset 2024) (48,944 spots for 18,946 genes, Fig. 1F). We first investigated how much metaspots impacted the computational resources needed to analyze spatial transcriptomic data (i.e., without considering the metaspot building step). For every γ from 1 to 25, we built metaspots with the procedure described in Figure 1A. We then performed normalization and computation of spatially variable genes with the Seurat (Hao *et al*. 2024) R package. We observed a rapid decrease of both time and memory usage with higher γ (Fig. 1G).

We further explored the metaspots obtained following the procedure in Figure 1B, i.e. after splitting based on cell-type annotations (final γ of 3.07) (Fig. 1F). The original gene expression matrix had 2.60% of non-zero entries while we reached 5.97% of non-zero entries within the metaspot gene expression matrix. This increase can also be observed by comparing the number of detected genes at the spot and metaspot level (Fig. 1H). In this dataset, the differences between spots and metaspots are mainly coming from the most abundant cell type (the Acinar.1 cell population, Supplementary Fig. 1A & B). Expression values of differentially expressed genes obtained at the spot level between the two most prevalent cell types, Acinar.1 and Ductal, show higher and more homogeneous expression for the marker genes at the metaspot level (Supplementary Fig. 1C). We further explored whether metaspots are compatible with standard visualization at the transcriptomic level. To this end, we compared UMAP analysis performed on the spots and metaspots (after SCT normalization (Hafemeister and Satija 2019) followed by PCA dimension reduction). We observed that cell types were equally distinguishable in both cases (Fig. 1I), suggesting that metaspots can be used for visualization of spatial transcriptomic data at the gene expression level.

Compared to metacells in scRNA-Seq data, the graining level that can be reached in metaspots is lower when imposing the constraint of purity with respect to cell type annotation. This is especially the case for tissues containing a high diversity of cell types within the same regions where the graining level is by essence limited by the spatial distribution of the different cells.

In summary, we extended the metacell concept to spatial transcriptomic datasets. Unlike approaches considering only spatial proximity to group spots (Morabito *et al*. 2023), our metaspots show promise in maintaining spatial and phenotypic information and improve computational efficiency for downstream analyses of spatial transcriptomic data. One limitation comes with spatially heterogeneous tissues where the constraints of spatial proximity and cell type annotation purity result in small graining levels. As such, we anticipate that metaspots will be especially useful for tissues containing relatively homogeneous regions, or technologies reaching sub-cellular resolution.

## Acknowledgements

We thank Yan Liu, Dana Moreno and Daniel M Tadros for testing the SuperSpot R package.

## Data availability

The data underlying this article is coming from publicly available resources. The 10x Visium Mouse Cortex raw data is accessible from the Globus endpoint ‘jhpce#HumanPilot10x’ (http://research.libd.org/globus) and the processed data from SpatialLIBD (Maynard *et al*. 2021) R package and website (https://research.libd.org/spatialLIBD/index.html). Nanostring CosMx Human Pancreas dataset and corresponding are directly available from Nanostring’s website (https://nanostring.com/products/cosmx-spatial-molecular-imager/ffpe-dataset/cosmx-smi-human-pancreas-ffpe-dataset/).

---

**Supplementary Figure 1:**
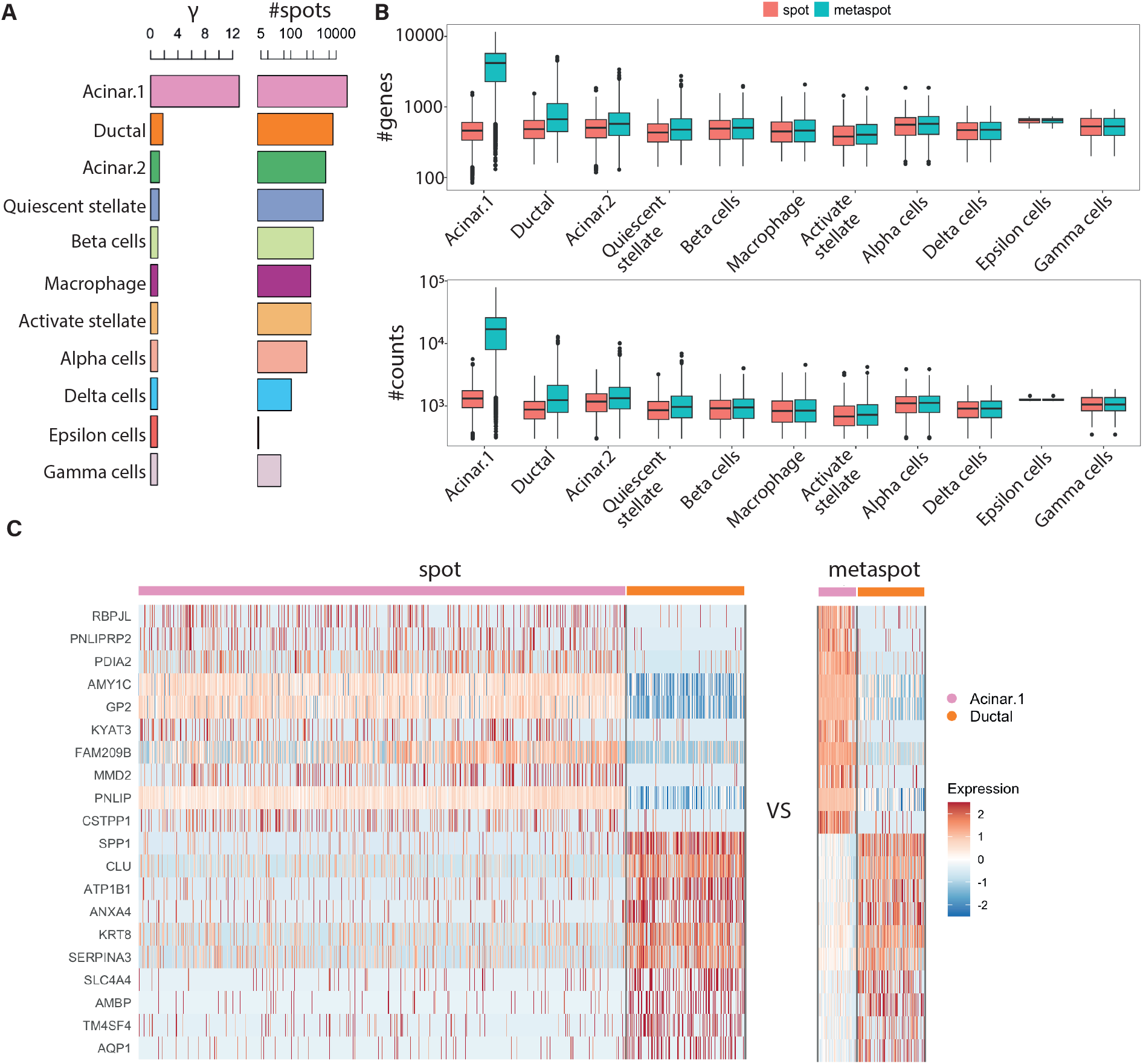
Analysis of metaspots across different cell populations in the Nanostring CosMx human pancreas dataset. **A)** Bar plots of the resulting γ and number of spots for each cell type. **B)** Boxplots of the number of genes and counts per spot and metaspot for each cell type. **C)** Heatmap of the expression at the spot and metaspot levels of the top 10 differentially expressed genes identified at the spot level between Acinar.1 and Ductal.

